# Chytrid fungal infections in alpine toads proliferate during winter dormancy

**DOI:** 10.1101/2025.07.07.663599

**Authors:** David R Daversa, Rob Grasso, Molly Posta, James O. Lloyd-Smith, H. Bradley Shaffer

## Abstract

The spread and impact of wildlife pathogens is often seasonal, and identifying the seasons of high impact is critical to biodiversity and public health management. Here, we report new evidence that winter host dormancy, a period generally neglected in terms of pathogen seasonal dynamics, promotes the spread of infection from the globally threatening amphibian fungal pathogen, *Batrachochytrium dendrobatidis (Bd)*. *Bd* surveillance of Yosemite toads (*Anaxyrus canorus*) during their first year of life showed that: (a) detectable *Bd* infections were nearly absent in the tadpole stage and immediately after metamorphosis, (b) *Bd* prevalence and intensity gradually increased during the first two months of post-metamorphic growth, and (c) *Bd* prevalence and intensity increased four-fold in toads during the first winter dormancy. High prevalence and intensity of *Bd* infections after winter dormancy was observed in yearling toads across two consecutive years and was much higher than that observed in aquatically breeding adults during the same timeframe. To our knowledge, this is the first evidence from wild amphibians that *Bd* proliferates during terrestrial winter dormancy. This discovery calls for a reconsideration of the seasons enabling the proliferation and persistence of *Bd* and identifies recently metamorphosed hosts as overwinter *Bd* reservoirs. More generally, the study underscores the importance of host dormancy in pathogen persistence and seasonal infection spread.

## Introduction

Many species survive in seasonally extreme environments by entering dormancy [1,2]. Hibernation in mammals, brumation in amphibians and reptiles, and aestivation in invertebrates (reviewed by [2]) are independently derived examples of this widespread phenomenon. Although dormancy is a widespread and well-studied life history strategy, the relationships between host seasonal dormancy and pathogen spread have been largely neglected [3]. The few host-pathogen systems in which host dormancy has been studied warn that this period may by key to the persistence, spread, and virulence of infections [4,5]. The clearest warning comes from lethal outbreaks of white nose syndrome (WNS) in hibernating bats, a disease caused by the fungal parasite *Pseudogymnoascus destructans* that is responsible for recurrent mass-mortality events across the eastern US and Europe [5]. Because host dormancy is accompanied by dramatic physiological and environmental changes that affect key epidemiological parameters (rates of pathogen exposure, transmission, persistence and growth), many species exhibiting this life history strategy may be vulnerable to dormancy-related disease outbreaks.

Here, we report new evidence that host dormancy is strongly associated with spikes in prevalence and intensity (i.e. proliferation) of infection from the fungal amphibian parasite *Batrachochytrium dendrobatidis (Bd)*. *Bd* is a microscopic chytrid fungus that thrives in moist, cool environments [6,7] and infects the keratinized skin of amphibians via motile aquatic zoospores [8]. Infection from *Bd* can cause the disease chytridiomycosis, which is a major ongoing cause of amphibian species declines and extinctions globally [9,10].

Many amphibian species pass winter months within terrestrial refugia (e.g., burrows, rock piles, etc.) in brumation, a hibernation-like form of dormancy involving extreme torpor. Because infective *Bd* stages are waterborne, short-lived and sensitive to drying [8,11], the use of terrestrial habitats for dormancy may be unfavorable to within-host *Bd* growth and persistence, and some field and laboratory studies support this expectation [7,12,13]. However, if terrestrial refugia used for dormancy remains sufficiently moist, *Bd* conceivably may survive on infected hosts or in the refugia environment [11,14,15]. Little direct evidence of either scenario exists in the extensive *Bd* literature. Yet, what is clearly evident is that *Bd* persists in many amphibian populations that have prolonged seasons of terrestrial dormancy, through at yet unknown pathways.

We carried out systematic *Bd* surveys in Yosemite toads (*Anaxyrus canorus*) to gain a deeper understanding of *Bd* dynamics in a host species with prolonged terrestrial dormancy. Yosemite toads are federally listed as ‘threatened’ under the US Endangered Species Act and a California Species of Special Concern [16]. The species is only found in a restricted part of the Sierra Nevada of California, USA where it occupies high-elevation meadows (1,950– 3,500 m) with prolonged winters characterized by freezing temperatures and heavy snowpacks. Yosemite toads are dormant (in brumation) from approximately October – April. During this period, they shelter in underground terrestrial refugia [17,18].

*Bd* presently circulates in wild populations of Yosemite toads [19], but the seasonal infection dynamics and conservation risk posed by *Bd* remain unresolved – a knowledge gap that has hindered population recovery initiatives. To address this gap, we carried out two systematic surveys across three years, six alpine meadows in Yosemite National Park, and all toad life stages (Fig. 1). First, repeated surveys of recently-metamorphosed toads (‘metamorphs’) throughout their first year of life characterized *Bd* dynamics before and after dormancy (Fig. 1c). Second, repeated surveys of all toad life stages – larvae, juveniles, and adult males and females - determined general life-stage differences in *Bd* infection prevalence and intensity throughout the active, non-dormant season (Fig. 1c). The survey results unambiguously identified the winter months of toad dormancy as a key season of rapid *Bd* proliferation, particularly in metamorphs. These results challenge most current thinking about the seasons of high *Bd* risk [20–22] and call for more concerted attention to infection dynamics during seasons of host dormancy.

**Fig. 1.**
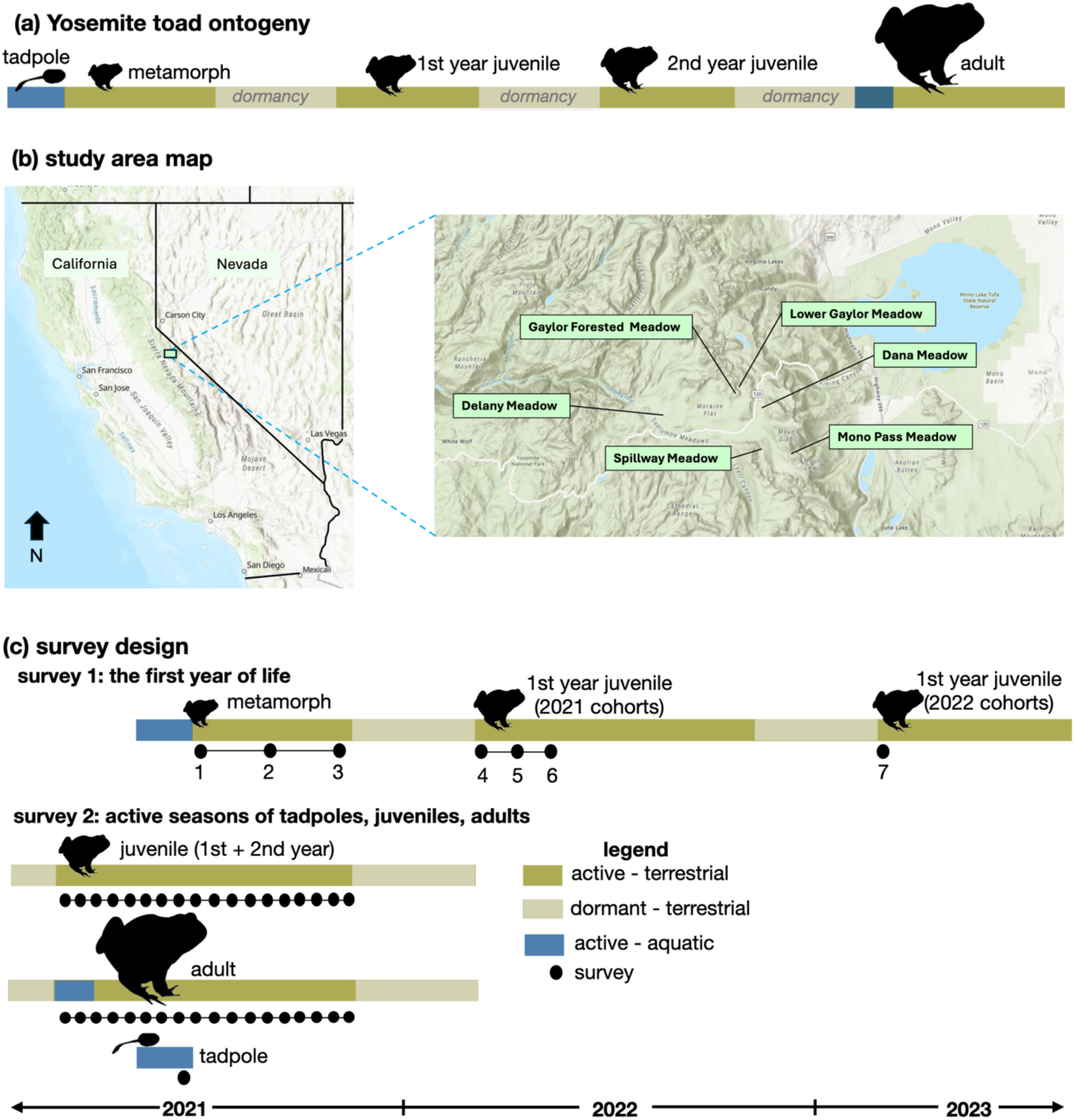
Yosemite toad (*Anaxyrus canorus*) ontogeny, study area map and survey designs. **(a)** Yosemite toads have a complex life history starting as fully aquatic larvae (tadpoles), which metamorphose into terrestrial ‘metamorphs’. Metamorphs emerge from their first dormancy as 1^st^ year juveniles, which then transition into larger 2^nd^ year juveniles following their second dormancy. Toads typically mature into breeding adults after the 3^rd^-4^th^ dormancy. **(b)** The six survey meadows in the Tioga Pass area of Yosemite National Park, CA, USA. **(c)** The designs of two repeated cross-sectional surveys beginning in 2021. The first survey followed toad cohorts throughout their first year of life, which included three monthly surveys (black dots) after metamorphosis and before the first dormancy, and three bi-weekly surveys (4-6) starting as soon as toads emerged from dormancy. We conducted a follow-up survey of 30 toads from the 2022 cohort as they emerged from their first dormancy in 2023.The second survey focused on tadpoles, juveniles, and adults during their active seasons and included one survey of tadpoles at Lower Gaylor meadow in the latter stage of development and 17 weekly surveys of juveniles and adults (black dots). The first two surveys of juveniles and adults were at the large breeding population at Lower Gaylor meadow. Surveys were expanded to the five other focal meadows from survey 3-17. In **(a, c)**, blue bars denote aquatic phases of the life cycle, including the larval stage and adult breeding seasons. Dark green bars are terrestrial active seasons (∼May – Oct), and light green bars denote dormancy, during which toads are dormant beneath snowpack in terrestrial underground refugia (∼Oct – April).

## Results

Toad metamorphs emerging at the same time and location formed distinct cohort groupings that persisted throughout their first year of life, which enabled repeat cross-sectional surveys of cohorts throughout pre- and post-dormancy periods. Surveying cohorts across six meadows in 2021 (Fig. 1b) revealed stark differences in *Bd* prevalence and intensity immediately before vs. immediately after winter dormancy (survey 3 vs. survey 4; prevalence: *X*^2^_1_ = 271.29, p < 0.001, Fig. 2a; intensity: *X*^2^_1_ = 479.92, p < 0.001; Fig. 2b). In the first survey after metamorphosis during mid-summer (survey 1), *Bd* was not detected at five of the six meadows (Fig. S1a). One month later, infections were detected in cohorts at all six meadows, and in general *Bd* prevalence (*X*^2^_1_ = 35.18, p < 0.001, Fig. 2a) and intensity (*X*^2^_1_ = 44.17, p < 0.001; Fig. 2b) increased in metamorphs over the first two months after metamorphosis. Across all meadows, *Bd* prevalence in metamorphs reached 23% (+/-5.5%) on average by the end of their first active season (Fig. 2a; survey 3).

**Fig 2.**
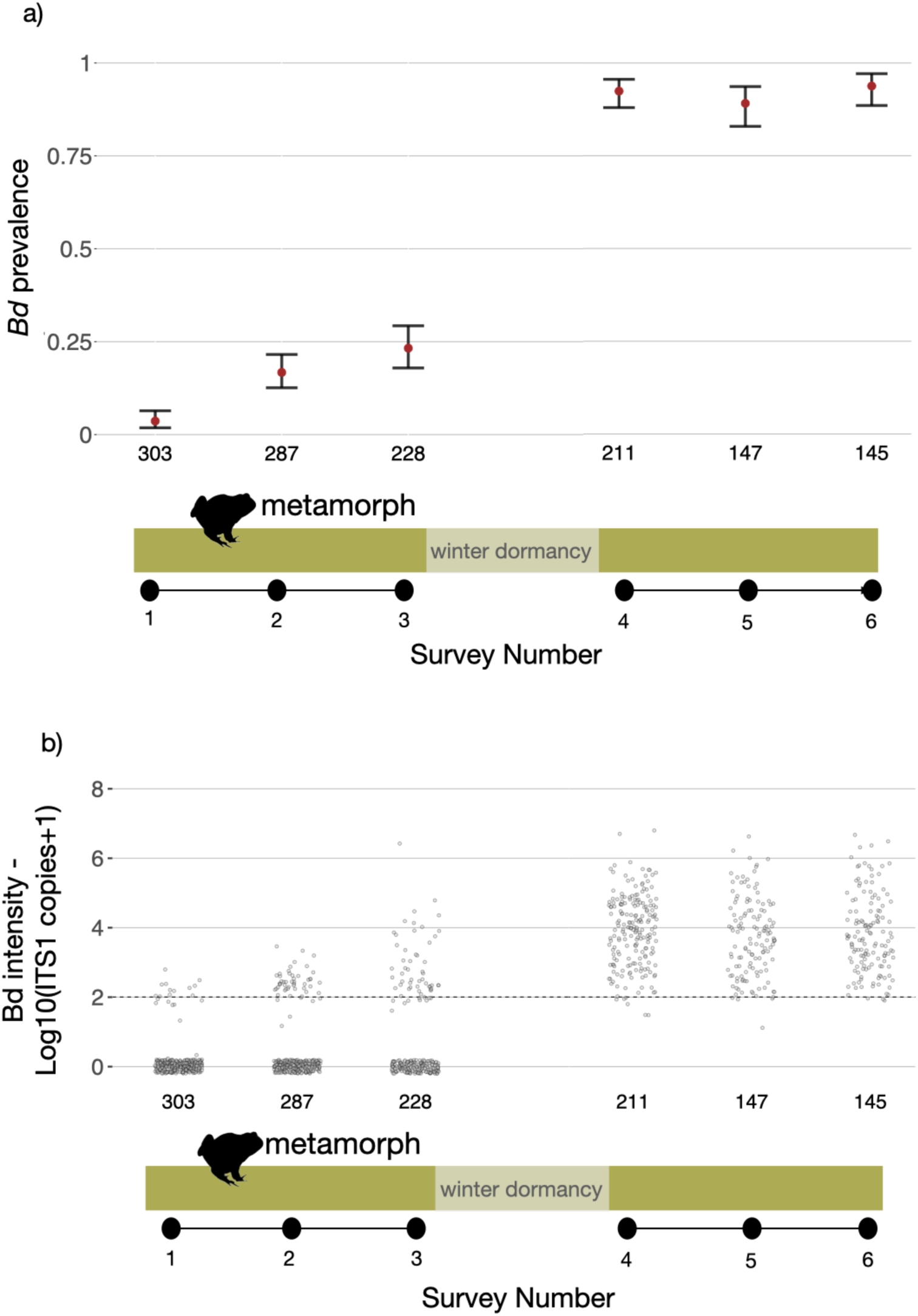
*Bd* dynamics in toad metamorphs before vs. after dormancy. Shown are **(a)** *Bd* prevalence and **(b)** *Bd* intensity observed in a longitudinal study of recently metamorphosed toads, which included six surveys and six different cohorts, each occupying a different alpine meadow. Survey 1 occurred immediately after metamorphosis, and subsequent samples were collected from the same cohorts one month (survey 2) and two months (survey 3) after metamorphosis in the same active season. We located the cohorts the following spring immediately after they emerged from dormancy and collected three additional samples every two weeks (surveys 4-6). In **(a)**, points denote overall *Bd* prevalence and error bars denote the 95% confidence intervals calculated using an exact test. In **(b)**, points represent *Bd* loads of sampled individuals, reported as the log-transformed number of IST1 copies (the gene region of *Bd* targeted by qPCR assays). We added 1 to the value to allow for log-transformation of zero values. The grey dashed line denotes the threshold ITS1 copy value for a clinical *Bd* infection (100, log-transformed +1). Sample sizes (number of individuals) are shown at the base of each plot.

The same cohorts exhibited >90% prevalence of infection when they emerged from dormancy the following year as first-year juveniles (Fig. 2a; Survey Number 4), and *Bd* prevalence remained high over four weeks of surveys (*X*^2^_1_ = 0.44, p = 0.505; Fig. 2a). Infection loads were also higher post-dormancy and remained at similar intensities in subsequent sampling (*X*^2^_1_ = 0.63, p = 0.427; Fig 2b). Parallel patterns were observed at all sites (Fig. S1). Eighteen individuals from a single cohort at Dana Meadow exhibited *Bd* loads exceeding 600,000 ITS1 copies upon emerging from their first dormancy (sample 4), loads which are generally associated with chytridiomycosis onset and mortality in Yosemite toad metamorphs (A. Lindauer, unpublished data) and sympatric host species [23]. No individuals showed overt signs of morbidity, however. To corroborate the above findings, we executed a seventh survey in 2023 of 30 toads immediately after their first dormancy (i.e. first-year juveniles of the 2022 cohort). There was a similarly high prevalence of infection (28/30 tested *Bd*-positive), with three individuals exhibiting *Bd* loads exceeding 600,000 ITS1 copies (Table S3), again without overt signs of morbidity.

The high prevalence and intensity of *Bd* infection in toads emerging from their first dormancy was consistent with *Bd* surveys of all toad life stages in a central breeding meadow (Lower Gaylor meadow) in 2021 (Fig. 1c). These surveys also followed a repeat cross-sectional design and showed that *Bd* prevalence and intensity differed across toad life stages in the first two weeks of the active season (prevalence: *X*^2^_3_ = 28.25, p < 0.001; intensity: *F*_3_ = 11.85, p < 0.001, Fig. 3). Although adult males and females were aquatic during this time, *Bd* prevalence and intensity was substantially lower in adult males and females compared to first- and second-year juveniles (Tukey’s tests; prevalence: z < 2.62, *p* < 0.044; intensity: *z* < 2.80, *p* < 0.031, Fig. 3). Adult males and females did not differ in early season *Bd* prevalence and intensity (Tukey’s tests; prevalence: *z* = 1.20, *p* = 0.630; intensity: *z* = 1.66, p = 0.353, Fig. 3). Early season *Bd* prevalence was highest in first-year juveniles recently emerged from their first dormancy, reaching up to 76.9%, though this was not statistically different from second-year juveniles, which exhibited 60% prevalence of infection (Tukey’s test: *z* = 1.23, *p* = 0.608, Fig. 3).

**Fig. 3.**
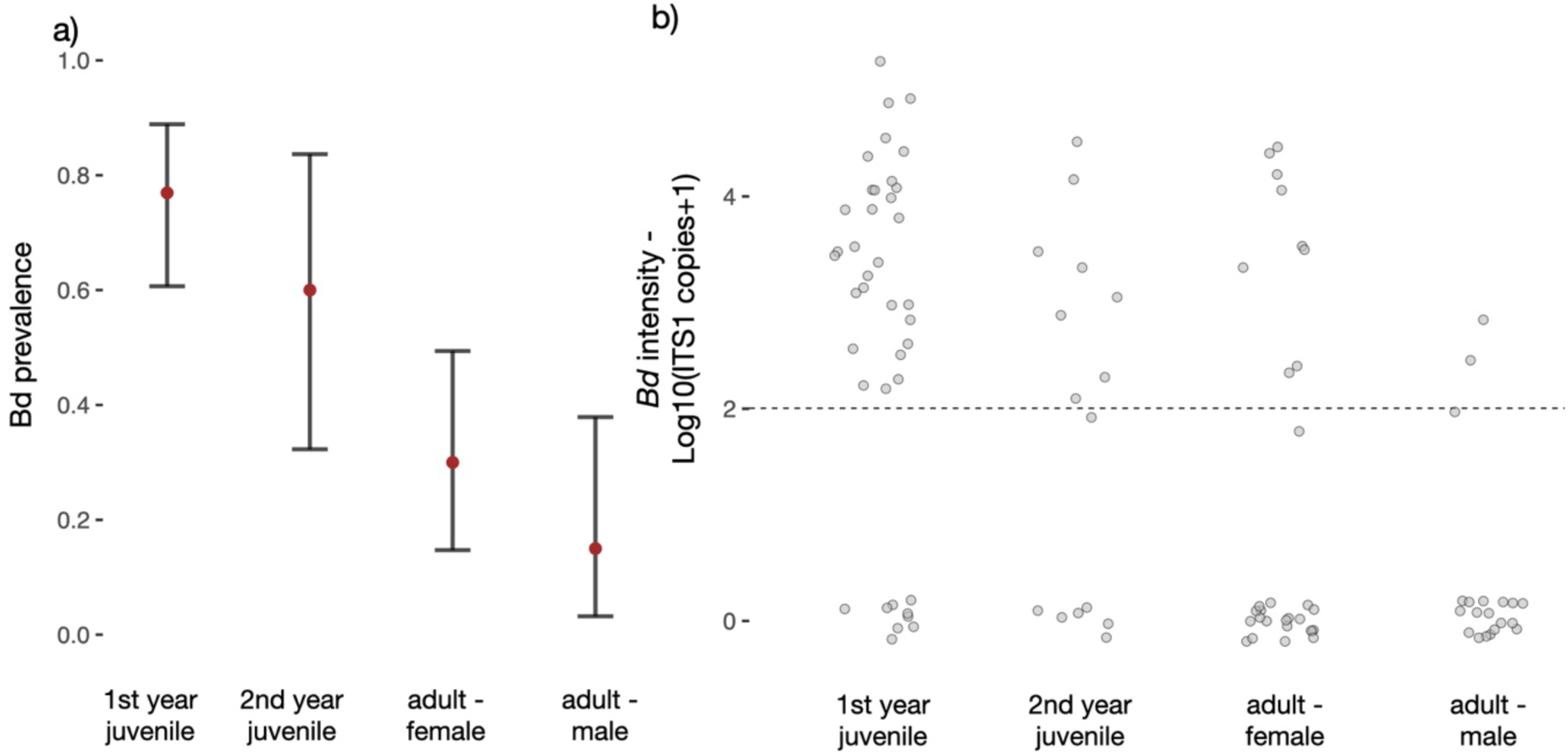
Early season *Bd* prevalence and intensity across toad life stages. The data presented are from the first two weeks of weekly *Bd* surveillance in post-metamorphic (metamorph, juvenile, adult male, adult female) Yosemite toads (*Anaxyrus canorus*) at Lower Gaylor meadow in Yosemite National Park, 2021. The surveys coincided with emergence from dormancy and adult breeding. **(a)** Points denote *Bd* prevalence and error bars denote the 95% confidence intervals estimated using an exact test. **(b)** Points denote the *Bd* infection intensity (i.e. load), measured as the number of ITS1 copies detected on skin swabs (+1 added to allow for transformation of zero values), and the grey dashed line denotes the threshold ITS1 copy value for a clinical *Bd* infection (100, log-transformed +1). M = male, F = female.

Continuing *Bd* surveillance in first- and second-year juveniles at Lower Gaylor meadow throughout the remainder of the active season showed that *Bd* infections were maintained at similar a prevalence. Broadening the survey of juveniles to five additional meadows in the Tioga Pass area similarly showed that *Bd* infections persisted well into the summer months of the active season (Fig. S2a). *Bd* infections were also detected in terrestrial adults during summer months at these meadows, but sample sizes were not adequate to reliably assess *Bd* persistence or temporal infection dynamics (Fig. S2b). No *Bd* infections were detected in toad larvae (*n* = 30) surveyed at Lower Gaylor meadow.

## Discussion

Pinpointing seasons of heightened infection risk is a key step toward effective disease control in wildlife [3,24], including amphibians [22,25]. Accomplishing this step for *Bd* has been hindered by seasonal dormancy exhibited by many host species, during which time hosts are often under snow or within inaccessible burrows, and thus difficult or impossible to survey. Our *Bd* surveillance in Yosemite toads indicates that more emphasis should be placed on *Bd* dynamics during host dormancy. Tracking metamorph cohorts before and after their first winter dormancy revealed that *Bd* prevalence was more than four-fold higher immediately after dormancy compared to immediately before, resulting in consistent peaks in *Bd* prevalence and intensity in the initial weeks of the active season across multiple, presumably independent sites and years. Although this infection peak coincided with adult breeding in aquatic habitats thought to be favorable to *Bd* growth and transmission [7,13,20,22], breeding adults exhibited fewer and weaker *Bd* infections than terrestrial juveniles. Taken together, these results suggest that *Bd* proliferates (increases in prevalence and intensity) during host dormancy, especially in early post-metamorphic life stages. To our knowledge, this is the first evidence from wild populations that *Bd* not only persists but also proliferates during terrestrial host dormancy.

Increasing infection prevalence and intensity during metamorph dormancy suggests a new intraspecific mechanism for interannual *Bd* persistence in host species that undergo winter terrestrial brumation. Current models invoke *Bd* resting stages [26,27], overwintering in larvae with extended larval periods [28,29], and alternative host species [22] to explain *Bd* persistence during dormant terrestrial periods in host annual cycles. Although such persistence mechanisms may be at play in certain host systems, our findings suggest a simpler, intraspecific model of *Bd* persistence in Yosemite toads in which external sources of infection need not be invoked.

The results of this study challenge current thinking regarding the seasons in host annual cycles that pose heightened *Bd* risk. In host species with life histories featuring terrestrial dormancy, *Bd* growth and transmission has been thought to be largely confined to active seasons, and specifically to activity periods that involve aquatic habitat use, such as adult breeding [20–22]. Infective *Bd* spores require moist environments [8], and negative associations between *Bd* risk (likelihood and intensity of infection) and host terrestrial habitat use have been demonstrated through experiments [7,30], field studies [12,13] and modelling [31]. The harsh winter weather of high-altitude meadows where Yosemite toad populations occur – with heavy snowfall and subfreezing air temperatures that fall well below the minimum temperature limit for *Bd* growth (∼2 degrees Celsius; Voyles et al. 2017) – further justifies the standard expectations of low infection risk during this part of toad annual cycles. Yet, the high post-dormancy prevalence and intensity of *Bd* suggests the opposite. In Yosemite toads, the risk of infection and/or infection growth is highest during winter dormancy, specifically in the first year of life. The environmental conditions of toad hibernacula must enable *Bd* to proliferate amid the hostile weather aboveground.

Laboratory studies with other amphibian host species indicate that *Bd* infections at metamorph life stages heighten mortality during dormancy [32,33]. Although we did not observe morbidity or mortality in metamorphs during our extensive field surveys, infected wild toads with high *Bd* loads certainly could have perished from chytridiomycosis without being detected. Experimental exposure of Yosemite toad metamorphs to high doses of *Bd* resulted in high rates of disease-induced mortality [34], demonstrating that *Bd* infections can be lethal to this life stage. Considering that the intensity of post-dormancy *Bd* infections in metamorphs approached levels known to be lethal in this and other host species [23], and that metamorphs are only 10-20mm in snout-to-vent length, cryptic *Bd*-induced mortality of wild metamorphs during dormancy is certainly plausible, and perhaps likely.

Pre-dormancy *Bd* dynamics in recently-metamorphosed toads during their first summer of growth also warrants further investigation. The lack of detectable *Bd* infections in tadpoles, which is consistent with previous work [19], indicates that hosts begin life *Bd*-free. Infections from *Bd* were also nearly absent immediately after metamorphosis, even though the keratinized skin cells that *Bd* infects should have then been present throughout the body [35]. Prevalence and intensity of *Bd* infection gradually increased during the first two months post-metamorphosis for unknown reasons. During this time, metamorphs gradually moved away from natal aquatic waterbodies into terrestrial (i.e., drier) upland habitat that should harbor less infective *Bd* stages, and so these patterns are not likely explained by increased *Bd* exposure. Although this initial increase in *Bd* prevalence and intensity was not pronounced, even this modest increase is puzzling and suggests that some infection intensification and transmission during terrestrial phases must be occurring.

Our findings establish the hypothesis that host dormancy can play an important role in driving transmission dynamics that enable population-level persistence of this often-lethal pathogen. To test this hypothesis, we encourage research that address three questions posed by this study. First, were post-dormancy spikes in *Bd* prevalence and intensity due to host-to-host transmission during dormancy, or alternatively due to growth of existing *Bd* infections to detectable levels [36]? Second, what factors – behavior [37], immunology [38], environmental conditions [27,39], or others – drive *Bd* proliferation during host dormancy? In a rare laboratory investigation of *Bd* dynamics during host dormancy, Kásler et al. (2023) found in juvenile agile frogs (*Rana dalmatina*) that *Bd* prevalence decreased throughout dormancy, in sharp contrast to our observations. Because frogs were housed individually by Kasler et al. (2023), these contrasting findings may indicate a role for host-to-host transmission driving *Bd* proliferation in dormant hosts.

Amphibians are known to aggregate in clumped groups in underground burrows during brumation [41], a behavior which we have observed in captive Yosemite toads, and this could be one explanation for our findings. Third, do *Bd* infections cause metamorph mortality during dormancy? Determining the extent of *Bd* transmission and mortality rates during metamorph dormancy, and the factors influencing these rates, would provide clearer insight into the population-level consequences of the *Bd* dynamics documented here.

The threats of fungal parasites extend to many wildlife that go through winter dormancy [42], and as such, our findings are likely generalizable to other systems. In addition to the well-known case of fungal *Pd* infections causing lethal White Nose Syndrome in bats, for which fungal proliferation during dormancy is well-established [5], other notable threats include the *Emydomyces testavorans* fungus that degrades the shells of turtles [43] and the *Ophidiomyces ophiodiicola* skin fungus that causes snake fungal disease [44]. All these pathogenic fungi have emerged as global threats in recent years and are a major focus of wildlife conservation. Like Yosemite toads, the hosts of these pathogens (snakes: Carpenter 1953, turtles: Ultsch 2006, bats: Bouma et al. 2010, Hoyt et al. 2021) undergo winter dormancy in terrestrial refugia, form aggregations during dormancy [45], and likely experience immunosuppression during dormancy [47], conditions which are amenable to fungal proliferation [5]. Broader investigation of fungal infection dynamics during host dormancy is likely to reveal general mechanisms shaping seasonal drivers of fungal infection and disease. As global environmental change alters the duration and environmental conditions of wildlife dormancy, such knowledge will be key to projecting the future of infectious disease and its impact on wildlife.

## Methods

### Study site

Our main surveys occurred from 2021-2022 and spanned six meadow complexes in the Tioga Pass region of Yosemite National Park, California, USA: Lower Gaylor meadow, Dana meadow, Delaney meadow, Gaylor Forested meadow, Mono Pass meadow, and Spillway meadow (Fig 1b). A follow-up survey post-dormancy in 2023 took place at Dana Meadow. The meadows are open, high-elevation grasslands surrounded by conifer forest and include aquatic breeding habitat (shallow ephemeral waterbodies) and terrestrial non-breeding habitat (willows, mammal burrows, rock piles) [17,18].

### Sample collection

Our cohort *Bd* surveys spanned the full active season of metamorphs in 2021 and the beginning of the active season in 2022 and 2023 when individuals were emerging from their first dormancy (May/June) (Fig. 1c). We hand-captured up to 50 toad metamorphs per meadow on three different occasions during their first active season in 2021 (before brumation): immediately after metamorphosis (survey 1, July), one month post-metamorphosis (sample 2, August), and two months post-metamorphosis (survey 3, September) (Fig. 1). The final survey in mid-September coincided with cooling temperatures and daylength shortening that precedes brumation (Fig S4, [41,48]). Upon capture, we collected a skin sample for *Bd* detection using a sterile swab (MWE 113 medical wire, UK) rubbed over the toad’s belly 15 times, 5 strokes on each thigh (total 10 thigh strokes per individual) and 5 strokes on each hind foot webbing (total 10 foot strokes per individual), following Vredenburg et al. (2010). For the very small individuals (<11m), we ran 30 strokes over the ventral side of the body because the thighs and feet webbing were too delicate and small to sample accurately.

We located the same cohorts as they were emerging from dormancy in 2022 as first-year juveniles (Fig. 1a,c). To minimize the possibility of detecting *Bd* infections contracted after dormancy, we scanned meadows for toads every 2-3 days before bare ground emerged from snowpack (May 17^th^, 2022, Fig. S5c). We did not observe any toads in our initial surveys, suggesting that our surveillance began when individuals were still dormant. Aerial imagery supports this case with evidence of extensive snowpack on our initial survey weeks (Fig. S5). We first observed metamorphs in Dana Meadow on 25 May 2022 as snowpack was starting to recede (Fig. S5d). As the cohorts emerged (late May-early June, depending on the site), we captured and swabbed individuals at three time points: immediately after emergence (survey 4, N = up to 50/meadow), two weeks after emergence (survey 5, N = 30/meadow) and four weeks after emergence (survey 6, N = 30/meadow). Cohorts could not be re-located in Delaney meadow in 2022, and so only five sites were sampled for surveys 4-6.

To compare early season infection across toad life stages, we executed weekly *Bd* surveillance at Lower Gaylor meadow in 2021 as soon as toads emerged from brumation (27 May 2021 – 16 Sept, 2021, Fig. 1c). We focused early season surveys at this meadow because it has consistently high abundances of breeding toads [49]. Again, we began scanning the meadow when snowpack was still fully present (Fig. S5a-b), and initial surveys did not detect any toad activity. We first observed toads at Lower Gaylor meadow on 25 May, 2021 when snowpack was receding (Fig. S5b) and began the surveys then. During surveys, we walked the perimeter of the meadow and captured and swabbed male and female adult toads that were not in amplexus as well as 1^st^ (2020 cohort) and 2^nd^-year juveniles. We used a cutoff of 23mm snout-to-vent length to distinguish 1^st^ year (<23mm) from 2^nd^ year (23mm or longer) juveniles. This cutoff was based on published literature [17] and our data from the cohort surveys (the largest 1^st^ year juvenile captured in the first two weeks post-dormancy in 2022 was 22mm SVL). We also collected and swabbed thirty tadpoles (Gosner stage 30-40) from Lower Gaylor meadow in June. Swabbing of tadpoles included 30 strokes of labial tooth rows and mandibles, which is the only keratinized structure of tadpoles, and therefore the only potential *Bd* infection site [23]. Starting in the third survey week, we expanded our surveillance to the five other focal meadows to assess temporal *Bd* dynamics more broadly.

### Sample processing

We screened swabs for *Bd* using standardized DNA extraction and qPCR procedures [50,51]. We quantified *Bd* DNA (the ITS1 gene region) on swabs using qPCR diagnostics, negative controls, and four concentration standards serving as positive controls (number of ITS1 copies: 100, 1,000, 10,000 100,000). Swabs were processed in singlicate due to the documented efficacy of *Bd* detection from single samples [23,52,53] and any ambiguous outputs were rerun. We defined *Bd* intensity (i.e. *Bd* load) as the number of ITS1 copies. A sample was considered *Bd*-positive when the ITS1 copy number was 100 or greater, matching our lowest-level, most sensitive positive controls.

### Data analysis

We used generalized linear models (GLMs) to evaluate the extent to which *Bd* prevalence and intensity varied by season and host life stage. For models of *Bd* prevalence, we used a binary infection status (0 = uninfected, 1 = infected) of individuals as the response variable and a binomial error structure. For models of *Bd* intensity, we used log-transformed *Bd* loads (+1 to allow for transformation of zero values, as we did not have enough power to analyze infection loads of infected individuals alone), and a Gaussian error structure.

To evaluate within-season *Bd* dynamics in the 2021 metamorph cohorts, we ran generalized linear mixed models (GLMMs) with meadow ID as a random effect to account for site-level variation and the survey number (1-3) as a fixed effect to measure changes over time. We used the same modelling structure to compare *Bd* prevalence and intensity in cohorts before vs. after dormancy. We compared infection data from the last cohort survey in 2021 (sample 3, Fig. 1c) with the infection data from the first cohort survey in 2022 (sample 4, Fig. 1c), using GLMMs with the survey number as a fixed effect and meadow ID as a random effect.

We used GLMs to compare early season *Bd* prevalence and intensity across life stages at Lower Gaylor meadow. We ran GLMs with life stage as a fixed effect with four levels: overwintered metamorph, juvenile, adult male, adult female.

For all analyses, we tested the influence of survey number and life stage on *Bd* prevalence and intensity by comparing the fit of models containing those factors to models omitting them, using likelihood ratio tests (MASS package in R) following either a chi-squared distribution in the case of GLMMs and binomial GLMs, or an *F* distribution for GLMs with Gaussian error structures. When a significant effect of a factor was detected, we examined differences between factor levels with Tukey’s HSD tests, using the emmeans function in R (emmeans package). Ninety-five percent confidence intervals for prevalence and mean intensity estimates were derived from an exact binomial test based on Clopper & Pearson [54], using the binom.test function in R.

## Acknowledgements

The authors acknowledge funding from the La Kretz Center for California Conservation Science at UCLA and the Yosemite Conservancy. All authors thank Alexa Lindauer for processing our samples; Carol Blanchette and staff at the Sierra Nevada Aquatic Research Laboratory for providing incredible field accommodations; Roland Knapp, Thomas Smith, and the Mountain Lakes Research Group for facilitating fieldwork planning; Rachel Hallnan for assisting the acquisition of environmental data and aerial imagery.

## Author Contributions

DRD, RG, HBS, and JLS contributed equally to idea formulation and planning. DRD, MP and RG completed field surveys. DRD and JLS executed the data analysis. DRD wrote the initial draft of the manuscript, which received multiple rounds of revisions by HBS, JLS, and RG.

## Data Availability Statement

Data and R scripts will be published on Figshare upon acceptance of the manuscript

