## Supplementary Material for "Chytrid fungal infections in alpine toads proliferate during winter dormancy"

### General differences in *Bd* prevalence and intensity across life stages

*Methods:* We pooled the survey data from 2021 and ran Generalized Linear Mixed Models (GLMMs) to evaluate broad differences in *Bd* infection prevalence and intensity across toad life stages, disregarding temporal and spatial variation. For models of *Bd* prevalence, we used a binary response (infected vs. uninfected) and a binomial error structure for models of infection prevalence. For *Bd* intensity, we used log-transformed *Bd* loads as the response, reported as ITS1 copies (+1 to allow for transformation of zero values, as we did not have enough power to analyze infection loads of infected individuals alone), and a Gaussian error structure. Both GLMMs included meadow ID and sampling week as random effects to account for site-level and temporal effects. The full models included life stage as fixed effect, and we included separate levels for adult males and adult females (4 levels total: 1st year juveniles, 2nd year juveniles, adult males, adult females).

*Results:* After accounting for site-level and temporal effects, *Bd* prevalence and *Bd* loads differed across toad life stages (Prevalence: $X^2_3 = 77.00$, $p < 0.001$, Fig. S3a; Loads: $X^2_3 = 137.19$, $p < 0.001$, Fig S3b). First- and second-year juvenile toads exhibited higher *Bd* prevalence (*Tukey's z ratio* > 3.9, $p < 0.02$, Fig. S3a) and mean *Bd* loads than other life stages (*Tukey's t ratio* > 4.91, $p < 0.001$, Fig. S3b). Adult males and females did not differ in *Bd* prevalence (*Tukey's z ratio* = 0.53, $p < 0.984$, Fig. S3a) or intensity (*Tukey's z ratio* = 01.19, $p = 0.755$, Fig. S3b).

| Meadow | Survey | Year | Sample size | Bd-positives | *Bd* prevalence | Bd load -mean | Bd load - SE |
|---|---|---|---|---|---|---|---|
| Dana | 1 | 2021 | 50 | 0 | 0.00 | 0.00 | 0.00 |
| | 2 | 2021 | 51 | 4 | 0.08 | 10.84 | 5.67 |
| | 3 | 2021 | 54 | 12 | 0.22 | 509.31 | 263.10 |
| | 4 | 2022 | 41 | 41 | 1.00 | 424654.80 | 235742.90 |
| | 5 | 2022 | 27 | 27 | 1.00 | 348816.40 | 194003.30 |
| | 6 | 2022 | 30 | 30 | 1.00 | 124690.20 | 66722.59 |
| Delaney | 1 | 2021 | 51 | 0 | 0.00 | 0.00 | 0.01 |
| | 2 | 2021 | 32 | 3 | 0.09 | 16.91 | 10.02 |
| | 3 | 2021 | 10 | 1 | 0.10 | 556.06 | 548.46 |
| Gaylor Forested | 1 | 2021 | 49 | 0 | 0.00 | 0.00 | 0.00 |
| | 2 | 2021 | 51 | 0 | 0.00 | 0.25 | 0.25 |
| | 3 | 2021 | 12 | 0 | 0.00 | 0.00 | 0.00 |
| | 4 | 2022 | 17 | 9 | 0.53 | 26383.08 | 16072.06 |
| | 5 | 2022 | 30 | 26 | 0.87 | 64593.21 | 32343.94 |
| | 6 | 2022 | 25 | 23 | 0.92 | 296508.50 | 192347.10 |
| Lower Gaylor | 1 | 2021 | 51 | 0 | 0.00 | 0.00 | 0.00 |
| | 2 | 2021 | 51 | 5 | 0.10 | 19.64 | 9.52 |
| | 3 | 2021 | 51 | 6 | 0.12 | 122.59 | 75.82 |
| | 4 | 2022 | 51 | 48 | 0.94 | 17056.59 | 8007.27 |
| | 5 | 2022 | 30 | 28 | 0.93 | 17789.60 | 6459.27 |
| | 6 | 2022 | 30 | 30 | 1.00 | 189059.50 | 82131.72 |
| Mono Pass | 1 | 2021 | 51 | 0 | 0.00 | 0.00 | 0.00 |
| | 2 | 2021 | 51 | 27 | 0.53 | 228.01 | 47.96 |
| | 3 | 2021 | 51 | 5 | 0.10 | 143.54 | 94.80 |
| | 4 | 2022 | 52 | 48 | 0.92 | 32777.62 | 11240.62 |
| | 5 | 2022 | 30 | 20 | 0.67 | 10690.70 | 7941.52 |
| | 6 | 2022 | 30 | 25 | 0.83 | 17749.66 | 8106.21 |
| Spillway | 1 | 2021 | 51 | 11 | 0.22 | 57.95 | 14.37 |
| | 2 | 2021 | 51 | 9 | 0.18 | 93.10 | 43.78 |
| | 3 | 2021 | 50 | 29 | 0.58 | 64819.05 | 60536.87 |
| | 4 | 2022 | 50 | 49 | 0.98 | 61357.89 | 16955.94 |
| | 5 | 2022 | 30 | 30 | 1.00 | 69195.17 | 41558.97 |
| | 6 | 2022 | 30 | 28 | 0.93 | 164756.80 | 136180.30 |

**Table S1.** Summary statistics of cohort survey of Yosemite toad metamorphs spanning six high elevation meadows in Yosemite National Park, CA, USA. Surveys began when toads metamorphosed in 2021 and continued monthly through the end of their first active season (samples 1-3). We returned after the 2021-2022 winter and resurveyed the cohorts when they emerged from dormancy (samples 4-6). The time interval between samples 4-5 and 5-6 was two weeks. *Bd* load represents the intensity of infection, measured in the number of ITS1 gene copies of Bd detected from qPCR. SE = standard error.

| survey week | sex/lifestage | sample size | Bd-positves | prevalence | Bd load - mean | Bd load - SE |
|---|---|---|---|---|---|---|
| 1 | Adult - Female | 25 | 7.00 | 3038.53 | 1669.64 | 0.28 |
| | Adult- Male | 10 | 0.00 | 0.00 | 0.00 | 0.00 |
| | 1st year juvenile | 6 | 4.00 | 15476.05 | 7881.57 | 0.67 |
| 2 | Adult - Female | 5 | 2.00 | 2401.24 | 1541.89 | 0.40 |
| | Adult- Male | 10 | 3.00 | 96.52 | 54.90 | 0.30 |
| | 2nd year juvenile | 17 | 10.00 | 9076.97 | 5946.57 | 0.59 |
| | 1st year juvenile | 39 | 26.00 | 9595.32 | 4185.27 | 0.67 |
| 3 | Adult- Male | 2 | 0.00 | 0.00 | 0.00 | 0.00 |
| | juvenile | 15 | 11.00 | 334783.22 | 328069.86 | 0.73 |
| 4 | Adult - Female | 5 | 0.00 | 0.00 | 0.00 | 0.00 |
| | Adult- Male | 2 | 0.00 | 0.00 | 0.00 | 0.00 |
| | juvenile | 33 | 14.00 | 1290.56 | 613.81 | 0.42 |
| 5 | Adult - Female | 4 | 0.00 | 0.00 | 0.00 | 0.00 |
| | Adult- Male | 7 | 2.00 | 43.11 | 20.77 | 0.29 |
| | juvenile | 52 | 20.00 | 3614.48 | 1933.14 | 0.38 |
| 6 | Adult - Female | 2 | 1.00 | 53.05 | 53.05 | 0.50 |
| | Adult- Male | 1 | 0.00 | 0.00 | NA | 0.00 |
| | juvenile | 16 | 6.00 | 12748.41 | 9449.45 | 0.38 |
| 7 | Adult - Female | 6 | 1.00 | 4034.42 | 4034.42 | 0.17 |
| | Adult- Male | 2 | 0.00 | 0.00 | 0.00 | 0.00 |
| | juvenile | 36 | 21.00 | 17595.78 | 8717.16 | 0.58 |
| 8 | Adult - Female | 2 | 0.00 | 0.00 | 0.00 | 0.00 |
| | Adult- Male | 2 | 0.00 | 0.00 | 0.00 | 0.00 |
| | juvenile | 31 | 14.00 | 13397.57 | 7218.38 | 0.45 |
| 9 | Adult- Male | 3 | 1.00 | 136.47 | 136.47 | 0.33 |
| | juvenile | 32 | 15.00 | 84850.61 | 45367.25 | 0.47 |
| 10 | Adult - Female | 1 | 0.00 | 0.00 | NA | 0.00 |
| | Adult- Male | 1 | 0.00 | 0.00 | NA | 0.00 |
| | juvenile | 5 | 1.00 | 3726.00 | 3726.00 | 0.20 |
| 11 | Adult - Female | 2 | 0.00 | 0.00 | 0.00 | 0.00 |
| | Adult- Male | 9 | 3.00 | 148.90 | 103.48 | 0.33 |
| | juvenile | 24 | 11.00 | 159496.65 | 153200.02 | 0.46 |
| 12 | Adult - Female | 3 | 1.00 | 80.70 | 80.70 | 0.33 |
| | Adult- Male | 3 | 1.00 | 34.53 | 34.53 | 0.33 |
| | juvenile | 16 | 4.00 | 70572.46 | 70104.96 | 0.25 |
| 13 | Adult - Female | 2 | 0.00 | 0.00 | 0.00 | 0.00 |
| | Adult- Male | 1 | 0.00 | 0.00 | NA | 0.00 |
| | juvenile | 11 | 4.00 | 1958.08 | 1496.99 | 0.36 |
| 14 | Adult - Female | 3 | 2.00 | 121.13 | 68.66 | 0.67 |
| | Adult- Male | 3 | 1.00 | 43.20 | 43.20 | 0.33 |
| | juvenile | 5 | 2.00 | 2576.24 | 2539.08 | 0.40 |
| 15 | juvenile | 2 | 2.00 | 214.25 | 58.55 | 1.00 |
| 16 | Adult - Female | 1 | 0.00 | 0.00 | NA | 0.00 |
| | Adult- Male | 1 | 0.00 | 0.00 | NA | 0.00 |
| | juvenile | 2 | 0.00 | 0.00 | 0.00 | 0.00 |
| 17 | juvenile | 8 | 1.00 | 112.51 | 112.51 | 0.13 |

**Table S2.** Summary statistics of population-level *Bd* surveillance of different Yosemite toad life stages during their 17-week active season. We used 23mm snout-to-vent length as a morphological cutoff to distinguish $1^{st}$ and $2^{nd}$ year juveniles, and we only made this distinction for the first two weeks of the active season (distinguishing the two became more ambiguous as the growth season progressed). We surveyed Lower Gaylor meadow exclusively during weeks 1-2, and this period coincided with adult aquatic breeding. Data from weeks 3-17 are pooled across six meadows in the Tioga Pass area of Yosemite National Park, CA, USA: Lower Gaylor meadow, Dana Meadow, Mono Pass meadow, Spillway meadow, Delany meadow and Gaylor Forested meadow. *Bd* load represents the intensity of infection, measured in the number of ITS1 gene copies of *Bd* detected from qPCR. SE = standard error.

| | date | Bd load | Bd status |
|---|---|---|---|
| 1 | 11-Jul-23 | 1446255.8 | 1 |
| 2 | 11-Jul-23 | 761666.5 | 1 |
| 3 | 11-Jul-23 | 582724.1 | 1 |
| 4 | 11-Jul-23 | 371022.8 | 1 |
| 5 | 11-Jul-23 | 329717.7 | 1 |
| 6 | 11-Jul-23 | 157855.9 | 1 |
| 7 | 11-Jul-23 | 111373.6 | 1 |
| 8 | 11-Jul-23 | 111269.4 | 1 |
| 9 | 11-Jul-23 | 59064.7 | 1 |
| 10 | 11-Jul-23 | 57439.5 | 1 |
| 11 | 11-Jul-23 | 37042.3 | 1 |
| 12 | 11-Jul-23 | 31766.1 | 1 |
| 13 | 11-Jul-23 | 31720.3 | 1 |
| 14 | 11-Jul-23 | 29651.5 | 1 |
| 15 | 11-Jul-23 | 27169 | 1 |
| 16 | 11-Jul-23 | 26700.5 | 1 |
| 17 | 11-Jul-23 | 26447.9 | 1 |
| 18 | 11-Jul-23 | 15667.1 | 1 |
| 19 | 11-Jul-23 | 11919.4 | 1 |
| 20 | 11-Jul-23 | 11586.5 | 1 |
| 21 | 11-Jul-23 | 10503.6 | 1 |
| 22 | 11-Jul-23 | 9965.8 | 1 |
| 23 | 11-Jul-23 | 8193 | 1 |
| 24 | 11-Jul-23 | 6698.4 | 1 |
| 25 | 11-Jul-23 | 5589.2 | 1 |
| 26 | 11-Jul-23 | 4710.9 | 1 |
| 27 | 11-Jul-23 | 1456.5 | 1 |
| 28 | 11-Jul-23 | 1110.9 | 1 |
| 29 | 11-Jul-23 | 0 | 0 |
| 30 | 11-Jul-23 | 0 | 0 |

**Table S3.** The table shows qPCR data from samples collected from metamorphs at Dana Meadow immediately after emerging from dormancy in 2023. The survey did not start until July because the winter preceding the 2023 active season was longer and wetter than normal. Thus, in addition to acting as a validation of our 2021-2022 surveys, these data also assessed whether the high *Bd* prevalence and intensity we observed post-dormancy in metamorphs is robust to differences in dormancy duration. *Bd* load represents the intensity of infection, measured as the number of ITS1 copies detected from skin tissue samples. *Bd* status is a binary measure of infection (0= uninfected, 1= infected), using an ITS1 copy threshold of 100 for a clinical infection (consistent with the lowest positive control in qPCR).

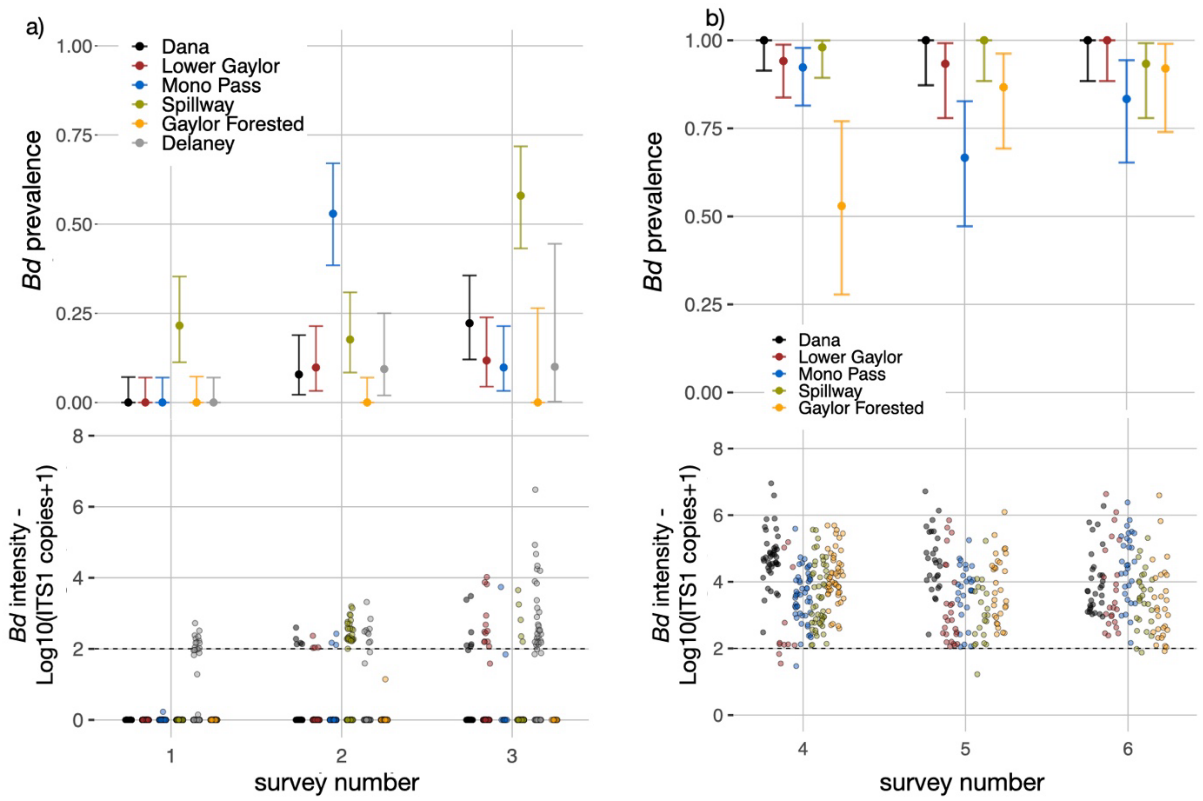


**Fig. S1. Site-specific *Bd* prevalence in toad metamorphs before vs. after dormancy**. *Bd* prevalence in toad metamorphs **(a)** from metamorphosis (sample 1) to time of entering their first dormancy in year 1 (samples 3) and **(b)** after their first dormancy in year 2 (samples 4-6). In the top panels, points denote the *Bd* prevalence at each meadow, and error bars denote the 95% confidence intervals calculated from exact tests. In the lower panels, grey dashed line denotes the threshold ITS1 value for a clinical *Bd* infection (100, log-transformed +1), with ITS1 copies being the amount of *Bd* DNA detected from skin swabs using qPCR. We surveyed six meadows in total. We omitted samples from Delaney meadow after dormancy in year 2 because the metamorph cohorts were not found. Samples 1-3 were collected monthly, and samples 4-6 were collected every two weeks.

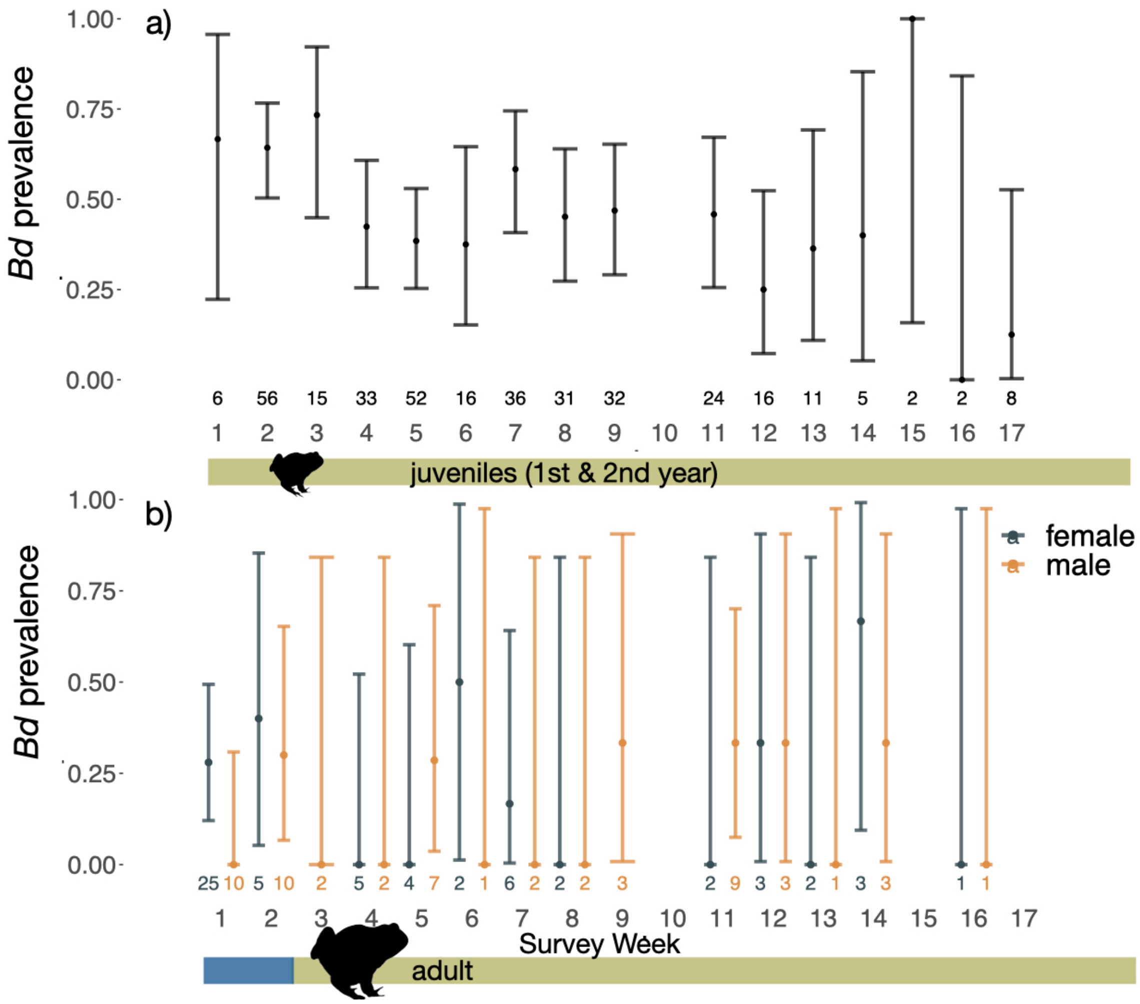


**Fig. S2. Temporal *Bd* dynamics over the course of the active season.** Infection data from the 2021 population-level survey of post-metamorphic life stages assessed *Bd* prevalence in **(a)** 1st and 2nd year juveniles and **(b)** adult males and females. Points denote the *Bd* prevalence and error bars denote the 95% confidence intervals calculated through an exact test. Weekly sample sizes are listed at the bottom of the plot. We surveyed Lower Gaylor meadow exclusively during weeks 1-2, and this period coincided with adult aquatic breeding. Data from weeks 3-17 are pooled across six meadows in the Tioga Pass area of Yosemite National Park, CA, USA. Sampling on week 10 did not occur due to inclement weather.

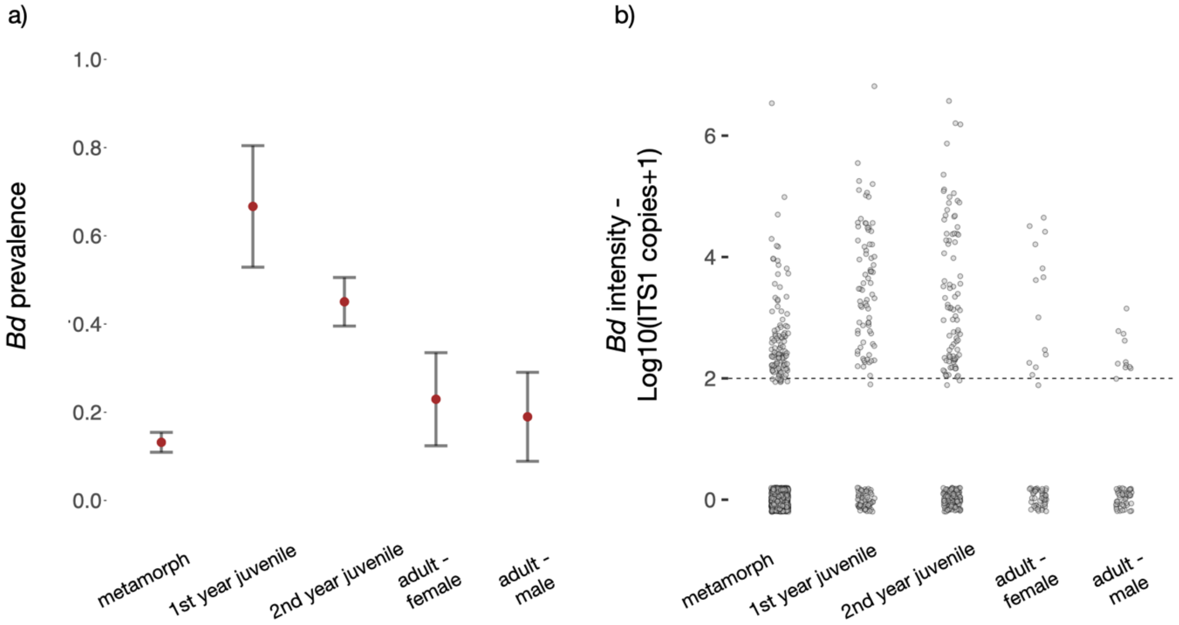


**Fig. S3. *Bd* prevalence and intensity across toad life stages.** Shown are the **(a)** prevalence and **(b)** intensity of *Bd* infections across all life stages of Yosemite toads sampled in 2021. Data were pooled across seventeen weekly surveys and six sites in the Tioga Pass area of Yosemite National Park, CA, USA. **(a)** Red points denote the mean *Bd* prevalence for a given life stage and error bars denote the 95% confidence intervals calculated from an exact test. **(b)** ITS1 copies denotes the amount of *Bd* DNA detected from qPCR of skin swabs, and the grey dashed line denotes the threshold for a clinical infection (100, log-transformed +1). First-year juveniles were presumably born and metamorphosed the previous year, and we distinguished them from 2$^{nd}$-year juveniles using a snout-vent length cutoff of 23mm. We considered all juveniles as 2$^{nd}$ years after the second week of surveys. Metamorphs emerged in July of 2021 (2022 data are omitted), half-way through the survey period, and were sampled monthly, for three months rather than weekly.

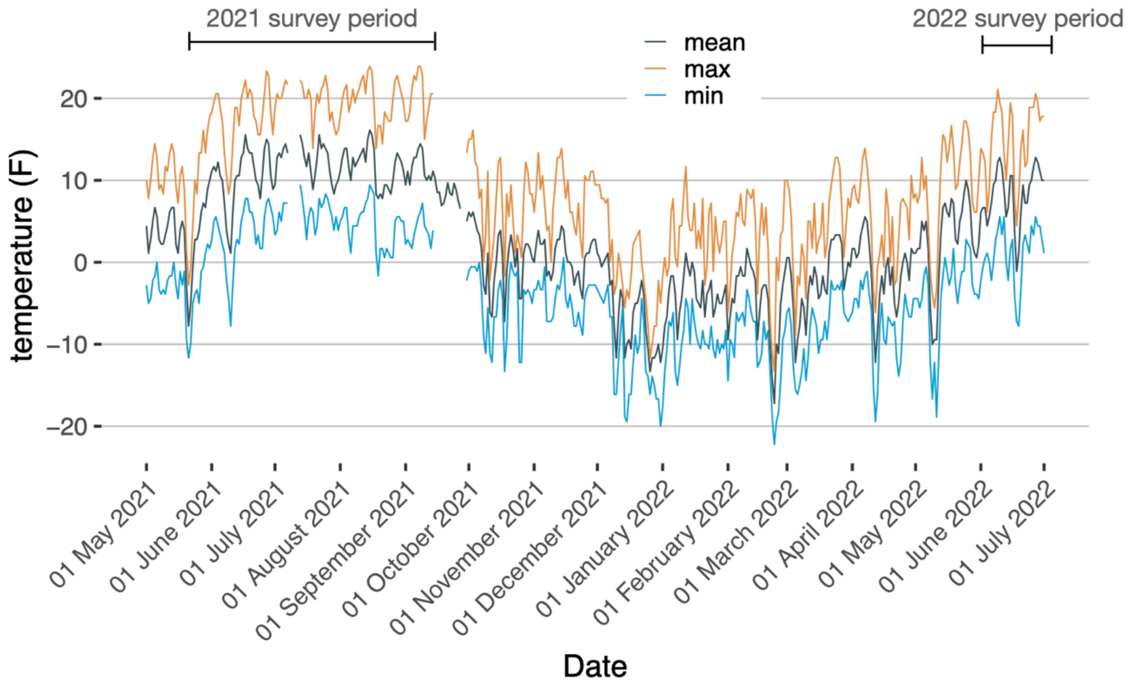


**Fig. S4.** Daily mean, maximum, and minimum temperatures at Dana meadow were recorded from a weather station operated by the California Department of Water Resources (https://cdec.water.ca.gov). Mean temperatures dropped in September 2021, when we ended toad surveys, which indicates the onset of dormancy, and rose in May of 2022 when we observed toads emerging from dormancy. Min and max temperature data were not available from 15-30 September, 2021, and mean temperature data were not available from 28-29 September 2021.

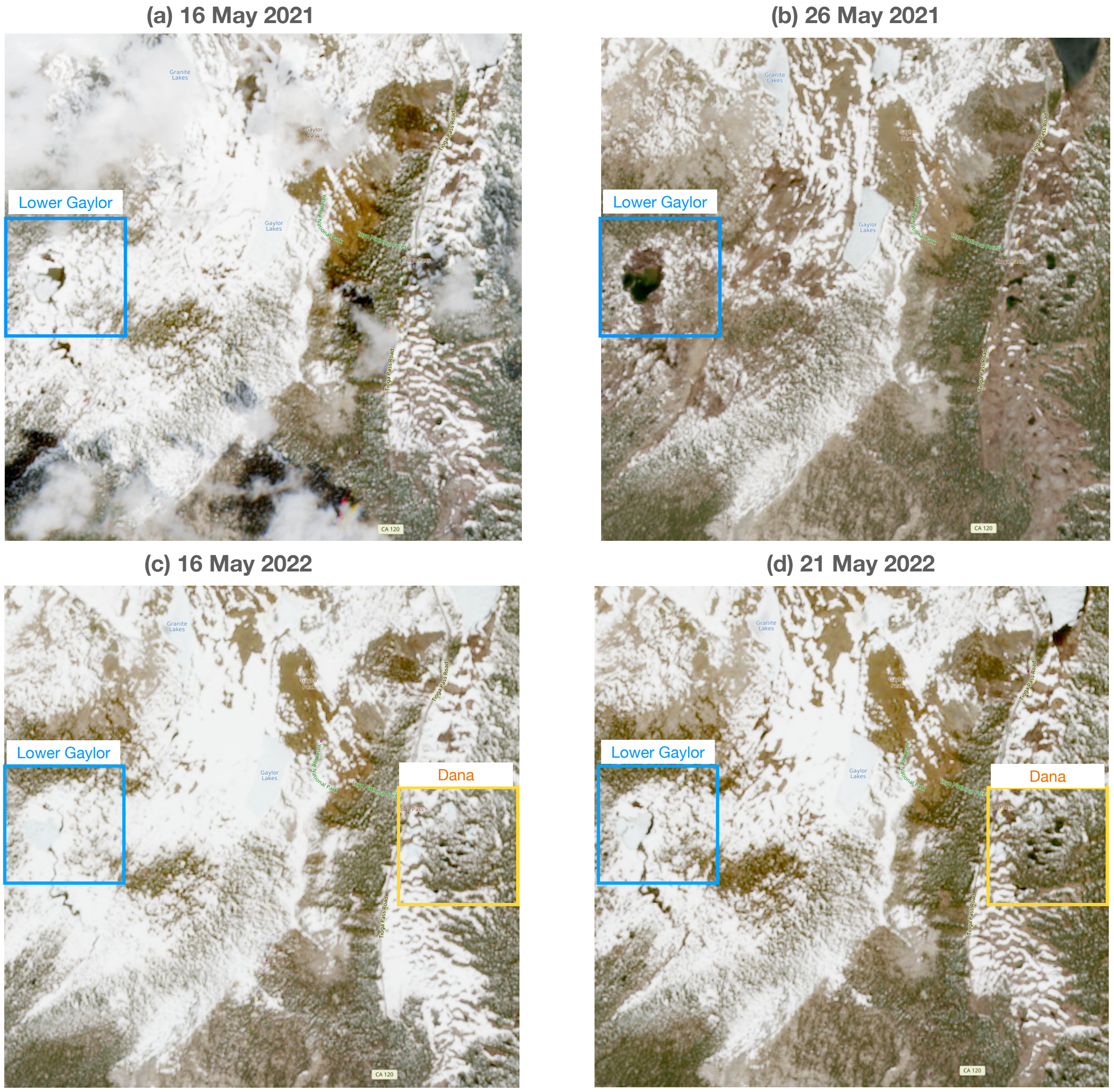


**Fig. S5.** Sentinel 2 aerial imagery shows snow cover for two focal meadows (Lower Gaylor – blue box; Dana – yellow box). The images show conditions within one week before the first toad survey of the season in **(a)** 2021 and **(c)** 2022, and within one-week after the first toad survey of the season in **(b)** 2021 and **(d)** 2022. On-site monitoring of toad activity began when snowpack was still present, and initial surveys did not detect any toad activity. In 2021, only Lower Gaylor meadow was monitored prior to toad emergence, and the first toads emerged on 25 May. In 2022, early season monitoring was extended the other five focal meadows, and the first toad emergences were observed at Dana meadow on 20 May. Imagery was access via the Copernicus online browser on 15 May 2025 (https://browser.dataspace.copernicus.eu).
